# Community conservatism is widespread throughout microbial phyla and environments

**DOI:** 10.1101/2024.11.06.622256

**Authors:** Lukas Malfertheiner, Janko Tackmann, João F Matias Rodrigues, Christian von Mering

## Abstract

Evolution gives rise to various long-term phenomena, including *Phylogenetic signal,* which describes the tendency of related biological taxa to resemble each other in morphology and function. Related taxa tend to also live in similar ecological niches – this trend is termed *Niche conservatism*. Both concepts are widely used to understand crucial aspects of evolu1on and specia1on and are well-established in animals and plants. The extension of these concepts to microorganisms is however challenging and thus far only assumed. Here, we hypothesize that two closely related microbial species, if indeed similar in both their morphology and their ecological niche, should be found in samples with similar community compositions. We propose “community conservatism” to refer to this, and leverage a database with millions of samples and hundreds of thousands of pairs of microbial taxa to assess their relatedness and the similarity of the communities they occupy.

Our findings reveal that community conservatism can be observed globally in all environments and phyla tested, to varying extents. Ecologically specialized taxa show a stronger community conservatism signal than generalists, and the signal can be discerned over nearly all taxonomic ranks. Analyzing community conservatism shows promise to advance our understanding of evolution, speciation, and the mechanisms governing community assembly in microorganisms. Furthermore, we propose it can be used to reintegrate ecological parameters into Operational Taxonomic Unit delimitation.

## Main

Phylogenetic lineages tend to retain their ancestral ecological niches over time ^1,2^. This so-called niche conservatism is often discussed in the context of a broader concept, phylogenetic signal, where closely related species tend to resemble each other morphologically and functionally ^3^. Numerous studies demonstrated niche conservatism and phylogenetic signal in animals and plants ^4–6^. Therein, the analysis and distribution of various traits, such as habitat preferences, morphology (e.g. leaf shape in Fig. 1A) and physiology, shed light on crucial aspects of evolution including speciation. Additionally, these studies help to predict how eukaryotes may adapt to rising challenges such as the spread of invasive species or climate change ^7,8^.

**Fig. 1:**
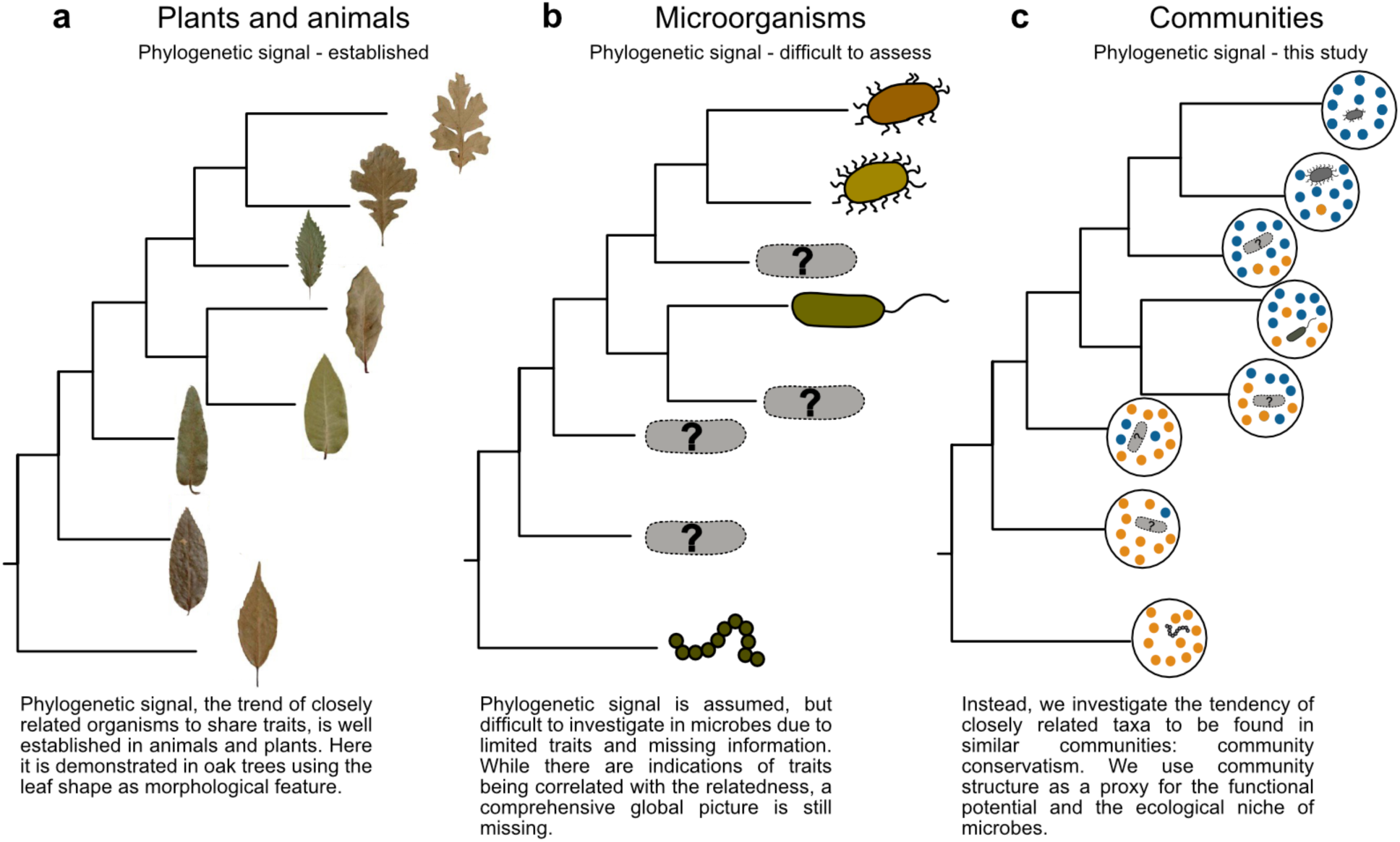
Community composition to measure evolutionary patterns in microbes. **a.** The leaf shape of oak trees is a morphological feature that shows a strong phylogenetic signal. Closely related species have similar leaf shapes, whereas more distantly related species have larger differences. Simplified/adapted from ref^6^. **b**. In bacteria, there are also indications that traits are phylogenetically conserved as in ref^17^. However, we often do not know enough about morphology or physiology of the organisms, since most of them remain uncultured. **c**. We propose community conservatism as an alternative approach: Instead of comparing the bacterial species directly in terms of physiology or morphology, we assume that if they are related (and thus have a similar function/occupy a similar ecological niche), also their community composition will be similar.

Apart from animals and plants, microorganisms also fulfill crucial roles in almost all areas of life, from driving biogeochemical cycles to influencing human health and diseases ^9–11^. Despite their importance, much less is known about the ecology and long-term evolution of microbes: only limited anecdotal evidence exists for niche conservatism and phylogenetic signal ^12–15^. For instance, studies indicate that some traits can be conserved over long time periods in microbes (Fig. 1b) ^16,17^. In computationally inferred microbial interaction networks, closely related organisms have been predicted to interact with one another more often (Phylogenetic assortativity) ^18–20^. However, most of these observations were restricted to a few environments and/or lineages.

The assumption that phylogenetic signal and niche conservatism are present globally is used in many popular algorithms, such as UniFrac and PINA ^21,22^. Characterizing microbial niche conservatism and phylogenetic signal on a global scale is thus crucial, yet challenging due to the lack of information about the traits of uncultured microbes ^10,23^. Additionally, microbes frequently transfer genes horizontally ^24^, which makes it difficult to infer traits from genomic information alone. The mix of vertical and horizontal inheritance furthermore impedes delineation of species boundaries and the assessment of evolutionary relationships ^25^. As a result, even the concept of species in microbes is still a matter of ongoing debate ^25–27^. Arguably, the currently used pragmatic method of delimitating Operational Taxonomic Units (OTUs) by solely considering sequence similarity at an approximate species-level threshold could also be improved by reintegrating ecological information.

Here, we look for an alternative to trait-based assessment of ecology, since environmental parameters are often not known and morphological features are scarce. Instead, we focus on what high-quality data we do have: millions of DNA-sequenced microbial community samples from all over the globe, and phylogenetic marker genes such as 16S rRNA enable us to know precisely in which communities a given microbial species occurs. Community structure can accurately distinguish different ecological niches ^28–31^ and has successfully been used to determine niche ranges in generalist and specialist animals and microbes ^32,33^. Additionally, communities are generally thought to be functionally relatively constant^34^.

Following this line of work, we here treat community composition as a proxy for the realized niche of a microorganism – the latter being determined through multiple, often unknown effects, ranging from the abiotic environment to microbial interactions. Thus, we hypothesized that by using a community-centric approach we can approximate phylogenetic signal and niche conservatism in microbes by analyzing the tendency of closely related organisms to occur in similar communities (Fig. 1c).

We show with an extensive analysis that more closely related taxa indeed occur in more similar communities. Remarkably, this trend is consistently detectable in all investigated phyla and environments. We suggest the term “community conservatism” for this phenomenon and show that remnants of microbial community preferences can be traced back billions of years. Furthermore, we show varying trends of community conservatism in different phyla, infer generalism- and specialism-specific signals and provide hundreds of OTU-pairs with potential interest for diverse research areas. Lastly, we outline the potential use of community conservatism as a second parameter – next to sequence similarity – in OTU clustering, to reintegrate ecological information in the future.

## Results and discussion

### Community structure as a proxy for niches and functional potential

We investigated global microbiomes utilizing the MicrobeAtlas project (https://www.microbeatlas.org), a database with a web-interface now containing over two million environmental microbiome sequencing samples, and over 400,000 taxa clustered into hierarchical OTUs using different similarity thresholds (from 90% to 99% full length 16S rRNA sequence similarity, whereas 97 to 99% traditionally correspond to “species-level” taxonomic groups^35^). By using such standardized occurrence data on a global scale, classical ecological questions can be investigated, such as the function of the ocean microbiome and which microbes are crucial for dissolving organic carbon ^30,36,37^.

We worked with OTUs defined at 99% sequence similarity (16S rRNA similarity) and initially assessed in which samples the OTUs occur globally (Fig. 2a). Next, we compared OTUs in a pairwise manner and calculated two main parameters for each pair: (i) the relatedness of the involved OTUs, and (ii) the similarity of the communities in which they occur (Fig. 2b). Relatedness is estimated from a large, high-quality phylogenetic tree, from which we randomly sampled pairs of OTUs to obtain a uniform distribution of phylogenetic distances (Extended Data Fig. 1). We then assessed the ß-diversity of all communities in which we detected them. For each pair, all samples containing the first OTU are compared to all samples containing the second OTU, measured as Bray-Curtis similarity (BCS: 1 - Bray-Curtis dissimilarity). Previous work showed that BCS can adequately distinguish ecological niches ^33^ and it can be computed at scale using optimized software ^38^. Lastly, pairwise plotting of relatedness and average community similarity values of each OTU-pair – combined with curve fitting – is used to assess the community conservatism signal (Fig. 2C).

**Fig. 2:**
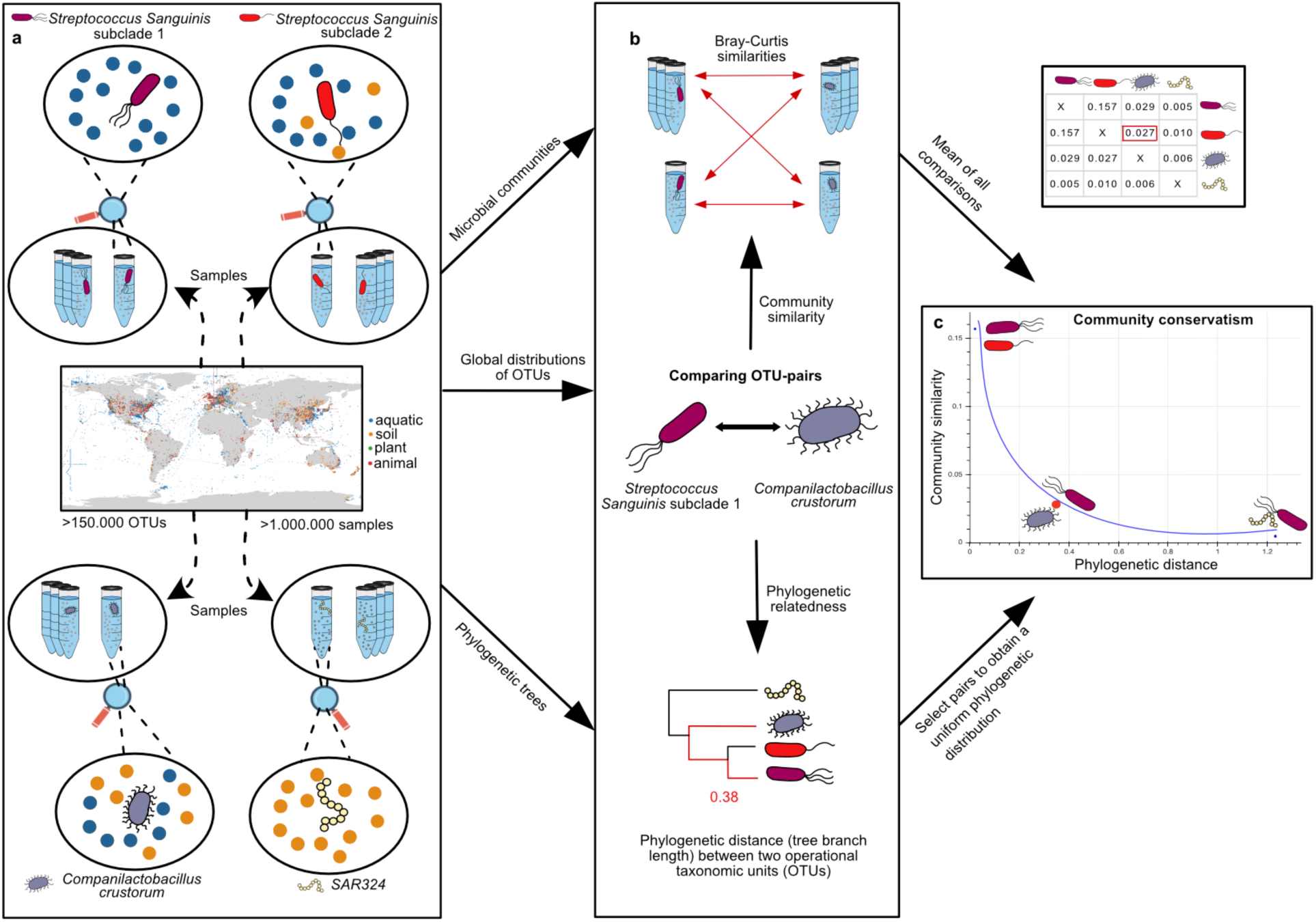
Analysis workflow. **a.** Illustration of the workflow using four selected example OTUs: Two closely related *Streptococcus sanguinis* subclades, *Companilactobacillus crustorus* still belonging to the same phylum (Bacillota) but a different family, and a completely unrelated *SAR324* strain. Within the MicrobeAtlas database all microbial sequencing samples (and their communities, respectively) matching strict quality filters are retained for testing, resulting in a global picture of the communities in which each OTU occurs. **b**. We compared OTU-pairs using two main parameters: Their relatedness, estimated by the tree branch length, and the average of all ß-diversity calculations from the communities in which they are found. **c**. After selecting test pairs following a uniform phylogenetic distribution, we visualize all selected pairs in a scatter plot. Each dot is one OTU-OTU pair, with their relatedness shown on the x-axis and the average similarity of their communities on the y-axis. Pairs which are closely related and show a large community conservatism are expected on the top left, and unrelated pairs with different communities on the bottom right.

To illustrate the general workflow, we compared four example OTUs with one another, at varying levels of relatedness: *Streptococcus sanguinis* subclade 1, *Streptococcus sanguinis* subclade 2, *Companilactobacillus crustorus* and *Sar324* (Fig. 2). The two *S. sanguinis* clades, belonging to the Bacillota, are closely related commensals found in the oral cavity of humans ^39^ with very similar communities (average BCS of 0.15). *C. crustorus* is a more distantly related Bacillota OTU found in diverse environments, including the human microbiome ^40,41^. Thus, despite sharing some community members (BCS 0.03), *C. crustorus* occupies different niches and appears to be more generalist. Lastly, *SAR324* is a predominately marine bacterium which is found in different layers of the ocean ^42^. It is only distantly related to the other OTUs, and as expected also its inhabited communities are very dissimilar (BCS 0.005). Hence, our hypothesis that more closely related OTU-pairs occur in similar communities is supported in this small example.

### Community conservatism is present on a global scale

To extend this workflow to a global scale, we chose 25,000 strictly quality-filtered, taxonomically annotated OTU pairs. We first assessed their sample-by-sample co-occurrence, showing that related species tend to occur more frequently in the same samples (Extended Data Fig. 2). While likely biologically relevant, this signal would compound our observations by inflating ß-diversity values when comparing identical samples. To mitigate this effect, we chose a conservative approach and only compared samples which do not belong to the same research project (i.e., do not share the same “project-ID” at the Sequence Read Archive).

We aimed to select the OTU-threshold which best reflects the ecological niche for the computation of ß-diversities. While we observed the same community conservatism trends with 90%, 97% or 99% OTU definitions (Fig. 3a, Extended Data Fig. 3), it has been hypothesized that microbial ecological niches are most clearly reflected at the genus level ^9,43^. In our dataset, 90% sequence similarity between OTUs roughly corresponded to a genus/family level divergence ^44^ (Extended Data Fig. 4). Additionally, more sequence reads can be unambiguously assigned when using 90% OTUs, thus we decided to use this level for all community similarity calculations going forward.

**Fig. 3:**
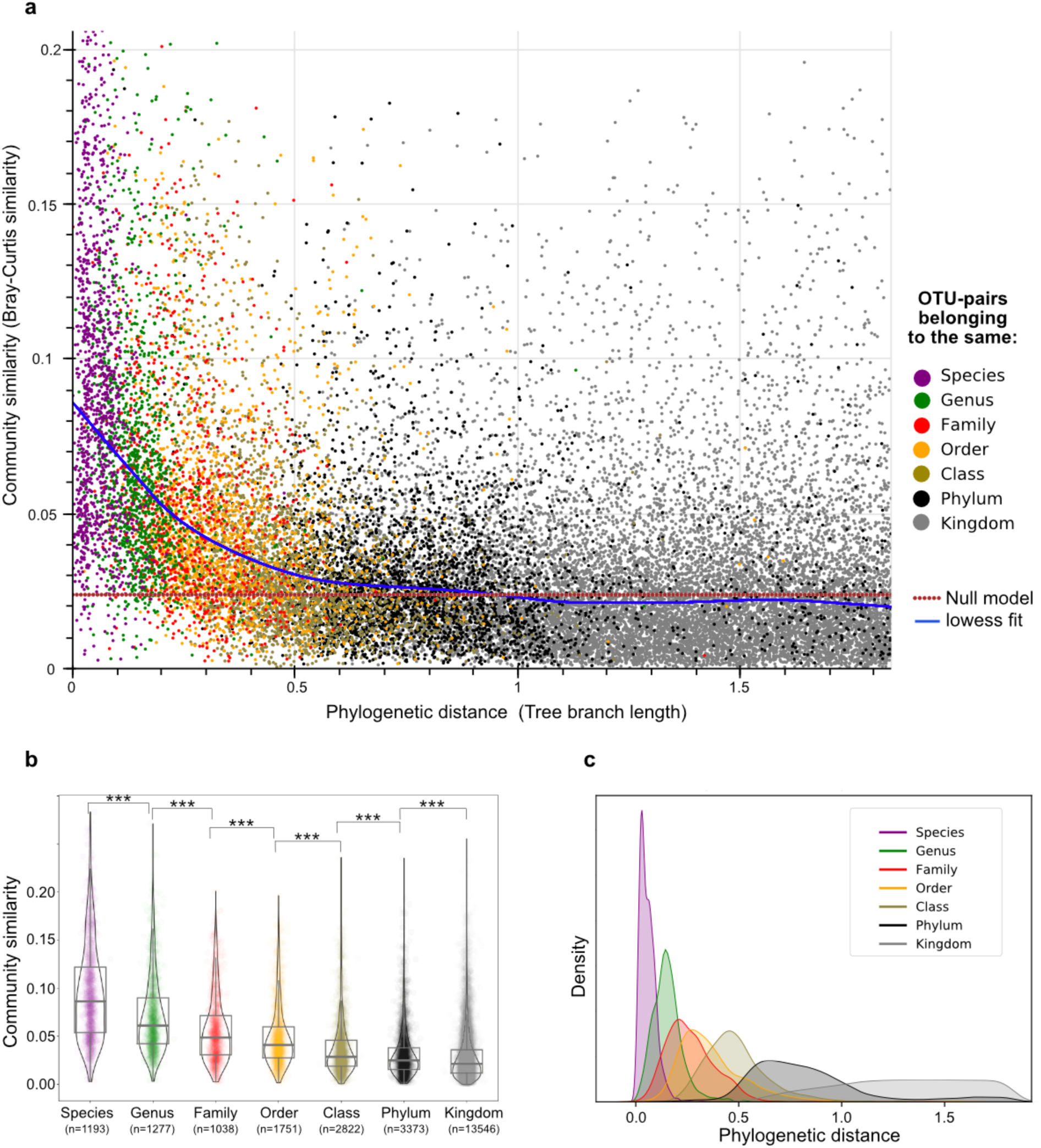
Community conservatism is present globally in microbes. **a.** Community similarity falls as phylogenetic distance increases, visualized here through 25,000 OTU pairs with available taxonomic annotation to species level; lowess fit and random expectation are shown as blue and red lines, respectively. Each dot corresponds to one OTU-pair colored according to their most specific shared taxonomic rank, with their relatedness shown on the x-axis and the average similarity of their communities (Bray-Curtis similarity) on the y-axis. **b**. All OTU-pairs are binned based on the most specific taxonomic rank they share. There are significant decreases in the community similarity-between all taxonomic levels, down to the phylum level (***: p_Mann-Whitney U_< 0.0001). Boxes correspond to mean +/− 1 standard deviation of each distribution. **c**. Density plot showing the phylogenetic distance distribution of pairs belonging to the same taxonomic groups.

Our results demonstrate the presence of community conservatism in microbes (Fig. 3a): OTU-pairs that are more closely related (x-axis, towards left) are more similar in their communities (y-axis, towards top). To visualize this observation, we fitted a locally weighted scatterplot smoothing (lowess, Fig. 3a) as well as an exponential decay function (Extended Data Fig. 5) to the data. We found both fitted curves strongly deviate from a null-model based on expected average community similarities between random samples (exponential decay coefficient: −4.23). Trends remain similar when using medians or other percentiles to aggregate community similarities (Extended Data Fig. 6).

To obtain a statistical estimation of community conservatism, OTU-pairs were furthermore binned according to their latest shared taxonomy. Significant deviation (p<0.0001) above the baseline was observed for each taxonomic level to the next, all the way up to phylum level (Fig. 3b). This indicates that community preference is traceable back – and has potentially been passed on – for billions of years ^45,46^. The largest differences in the average BCS exist between species and genus levels, suggesting that species-level adaptations are particularly important for community preferences.

We showed that OTU-pairs belonging to the same species are often found in very similar communities, and hence competition due to their overlapping niches might be expected. The coexistence of many direct competitors should not be feasible according to classical ecological models ^47–49^. While this conflicts our observation that closely related strains are also often co-occurring (Extended Data Fig. 2), there have been more observations showing said co-occurrence ^50,51^. Recent research has shown that horizontal gene transfer might alleviate the competition between related microbial species and allow the coexistence of many closely related competitors ^52^. However, competition or exclusion of closely related taxa at a small scale – which remains undetected in our 16S rRNA based analysis – cannot be excluded.

Next, we checked how well the taxonomic relatedness of OTUs (based on available NCBI annotations) overlapped with the tree branch lengths that we use to estimate relatedness. Overall, taxonomic ranks follow phylogenetic distances as expected (Fig. 3c). However, it is also apparent that in some cases taxonomic classifications and 16S rRNA sequence similarities do not fully agree (Extended Data Figure 7). This is consistent with known deviations between trait-based taxonomies and purely sequence-based clustering ^53^.

### Environmental preferences are entangled with community conservatism

Consistent with the concept of niche conservatism, we postulated that related OTUs would tend to inhabit similar environments. To test this hypothesis, we used MicrobeAtlas environmental annotations to compute per-environment OTU abundances. Next, we categorized each OTU into one of five primary environments: soil, animal, plant, marine, and freshwater. Our analysis revealed a consistent trend for related species to be found in the same main environment, partially driving the observed community conservatism trends (Fig. 4a). These findings suggest that broad-scale niche conservatism, the tendency of OTUs to remain in their primary environments, is also evident in microorganisms. While we utilized only broad habitat classifications due to the often-unreliable nature of publicly available metadata, previous observations of niche conservatism at a smaller scale ^14,54^ indeed indicate that, given sufficient high-quality data, this concept could extend to more specific ecological niches.

**Fig. 4:**
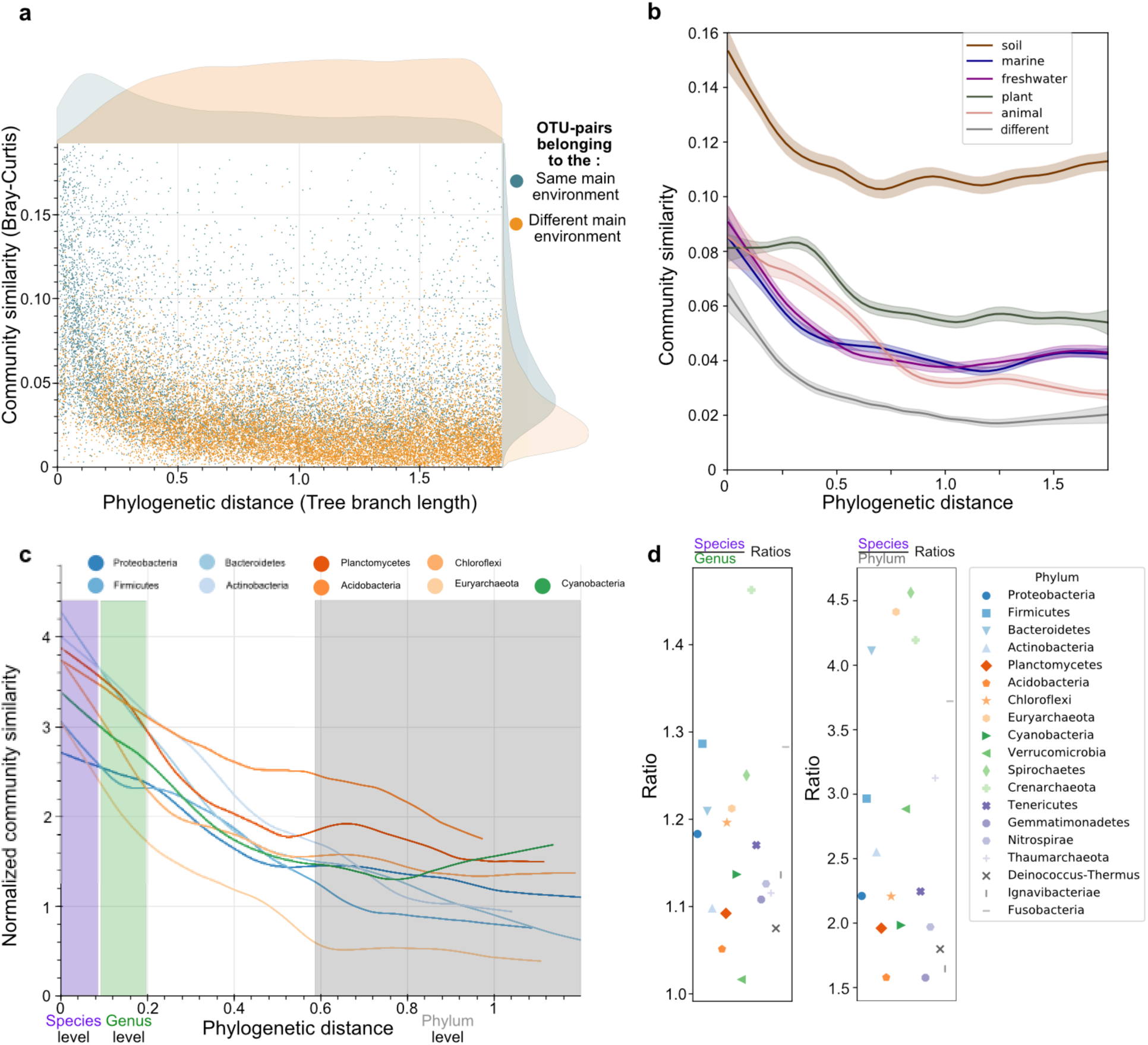
Environmental effects and phylum-level differences. **a.** The phylogenetic distance of 25,000 OTU-pairs is plotted against the similarity of the communities they occupy. The pairs are colored according to whether they share the same main annotated environment (blue) or are assigned to differing environments (orange). **b.** Only OTU-pairs belonging to the same given main environment (soil, marine, freshwater, soil or plant), or to different environments (grey) are compared. The curves calculated from their lowess fits (95% confidence interval) are plotted. **c.** All Phyla with >3,000 available OTUs are analyzed separately, and a lowess fit is created for each. The signal is normalized by environmental preference (see Methods). Additionally, taxonomic ranges (estimated from Fig. 3c) are indicated by color shade (purple=species, green= genus, grey=phylum) **d**. Community conservatism ratios calculated from the taxonomic bins (Fig. 4c) (left: species/genus; right: species/phylum) of all 19 investigated phyla.

We were furthermore curious whether community conservatism extends beyond these broad environmental preferences, i.e. whether it still persists even when the primary environment is normalized for. To investigate this, we limited the analysis to OTU pairs that were either assigned to the same main environment or to pairs belonging to different environments (Fig. 4b).

This analysis revealed that community conservatism is consistently observable even *within* different environments, interestingly with varying trends. Conversely, as expected, OTU-pairs annotated to different main environments had the lowest overall community similarities. Most environments exhibited similar BCS-ranges of their OTU-pairs, with the notable exception of soils, which showed almost twice the community similarity of other environments. While soil microbial communities can vary significantly even at centimeter scales, they are globally more similar than often assumed and are usually dominated by a few taxa ^55,56^. Additionally, most soil samples are agricultural and therefore particularly homogeneous. Interestingly, the community conservatism of OTUs mainly annotated to plants already plateaus at approximately genus-level phylogenetic similarity. This could potentially be rationalized by OTUs preferring certain plant types at a broader phylogenetic level. However, it is important to note that only 2,781 OTUs are predominately annotated to plants, which is 4-fold less than in any other environment (next lowest: freshwater 11,671 OTUs). This may increase the likelihood of artifacts stemming from a somewhat higher number of redundant comparisons across large phylogenetic groups.

### Phyla-specific characteristics of community conservatism

We showed that community conservatism is present in microorganisms on a global scale, irrespective of their main environment and extending as far back as the phylum level. The next question we wanted to address is whether we can infer characteristics of the ecology, speciation and community assembly processes across the different phyla. For this, we next repeated the previous analysis separately for each phylum represented by a minimum of 500 OTUs in the MicrobeAtlas database. In total, these were 3 Archaeal and 16 Bacterial phyla (Extended Data Fig. 8). As we showed previously, the environment in which the phyla are mainly found strongly influences community conservatism. To mitigate that effect, we calculated phylum-specific null models, considering the expected community similarity values by accounting for the main environments of the compared OTUs (Supplementary Table 1).

By normalizing our community similarity metrics against these phylum-specific baselines (see Methods), we obtained normalized community conservatism curves. Intriguingly, these curves trend differently across phyla, with those containing less than 3,000 OTUs showing increased noise (Extended Data Fig. 9). Yet, most phyla show a clear decrease in community conservation when assessing increasing taxonomic distances from species level to phylum level. We see a steep descent in some phyla (e.g. Crenarchaeota, Firmicutes), while in others the decrease is more gradual (Fig. 4c).

We hypothesized that quantifying the steepness of this trendline, as well as differences to the baseline, would help us to characterize ecological characteristics of each phylum. Since not all OTUs were taxonomically annotated to the species level, we instead used the density gradients obtained in Fig. 3c and binned the OTUs accordingly to approximate the taxonomic levels. In all investigated phyla we observed highly significant (p<0.0001) decreases in community conservatism when comparing the species level to the phylum level. We furthermore quantified the decrease of community conservatism from species level to genus level (Fig. 4d). We reasoned that for phyla with only early, conserved community preferences which are not currently changing, comparison at the species and genus level should yield similar conservatism signals. However, we found 15 phyla that were still significantly different in their community conservation when comparing species level versus genus level, suggesting ongoing changes in community preferences (Supplementary table 2). For instance, Crenarchaeota have large increases at both levels (Species/Genus Ratio 1.47, Species/Phylum ratio: 4.25), showing a strong and ongoing tendency to change communities and to specialize into different niches. But all investigated archaeal phyla show sharp increases (species/phylum ratios > 3), aligning with their tendency to be found in extreme environments and their adaptability to new environmental factors ^57^. On the other hand, phyla such as Acidobacteria, predominately found in soils ^58^, show a comparatively shallow increase (Species/Genus Ratio 1.05, Species/Phylum ratio: 1.6). This indicates that members of this phylum have long been restricted to their respective niches and do not usually adapt and evolve quickly into new habitats or roles.

This examination provides a quantifiable measure of similarity among related OTUs, a concept central to methods like UniFrac. UniFrac compares communities by considering phylogenetic relationships (i.e., closely related species are assumed to be similar and thus contribute less to diversity) through tree branch length calculations^21^. However, UniFrac defines relatedness for all microbes equally, while our study reveals that different phyla exhibit varying rates of community similarity with decreasing relatedness. We propose that the values presented in our analysis could be utilized to develop an ecologically informed version of UniFrac in the future. This enhanced method would apply phylum-specific weights when calculating tree branch lengths, potentially offering a more nuanced approach to community comparison.

### Specialists and generalists have distinguishable community conservatism trends

Conceivably, the observed differences between phyla in terms of community conservatism might hint at general differences in their degree of ecological specialization: When members of a phylum show little specialization (i.e., they are generalists) we would expect their communities to be fairly diverse, with community-community distances averaging out at certain level set by the overall diversity of the available data. Conversely, in phyla predominantly composed of specialists, closely related pairs would be hypothesized to share very similar communities, whereas the communities of pairs with a larger phylogenetic distance between them are expected to be very dissimilar due to very distinct niches.

To check for this, we first devised a habitat generalism score for each OTU, based on their normalized abundances across different environments (see Methods). We then selected the top 10% OTUs (“Generalists”), and the bottom 10% OTUs (“Specialists”), and calculated the community conservatism of both groups. Strikingly, the results reveal a clear separation: Generalists show small, steady increase of community conservatism, with a relatively high baseline even in non-related pairs, whereas specialists show a much steeper trendline, with non-related pairs found in very different communities, whereas closely related pairs appear in very similar communities (Fig. 5a).

**Fig. 5:**
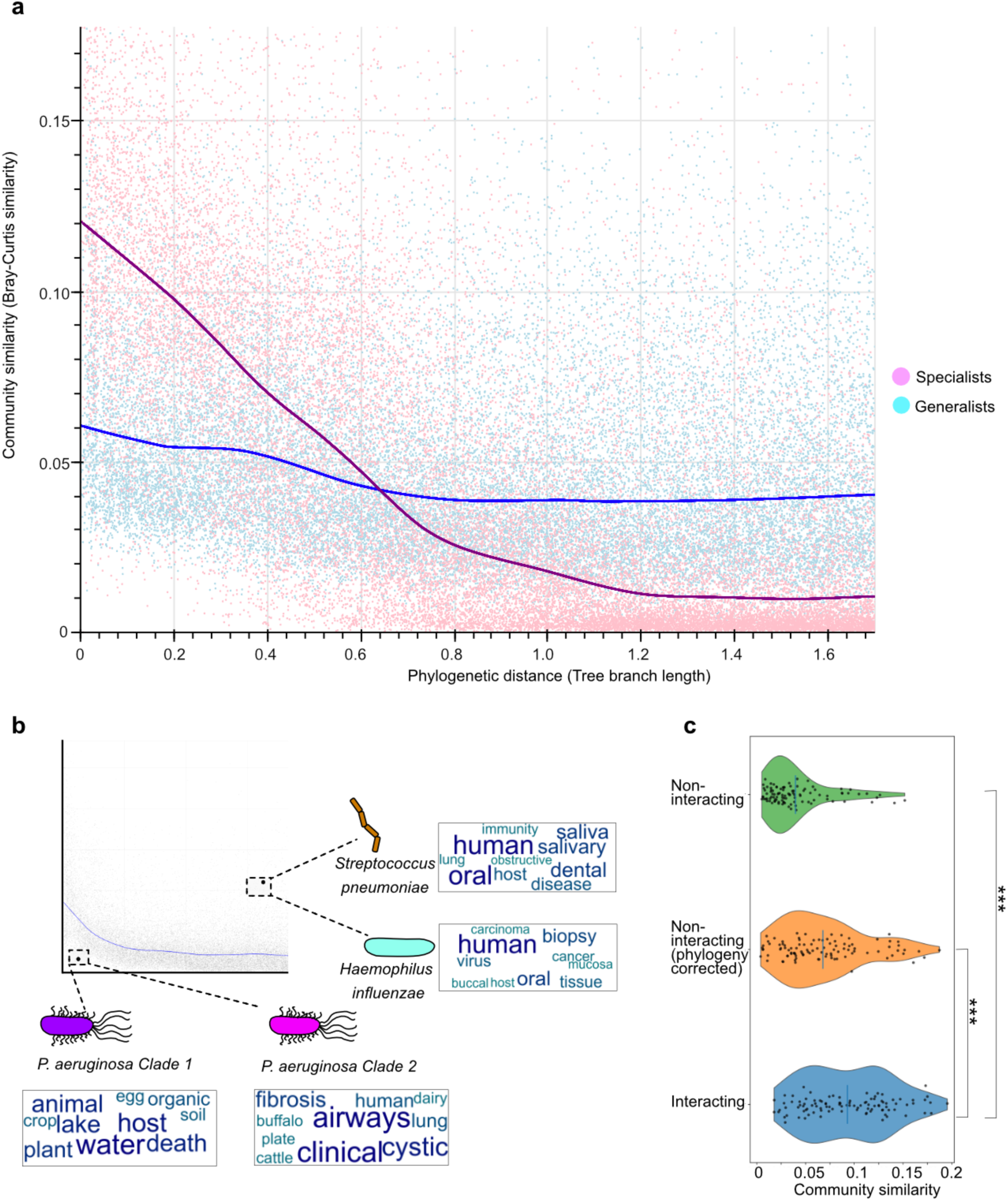
Community conservatism correlates with ecological properties. **a.** 10,000 OTU pairs consisting of only generalists (light blue dots) are compared to 10,000 specialist pairs (pink dots). Lowess fits of both groups (pink and light blue trendlines) are plotted on top. **b**. Two outlier OTU-pairs are highlighted here: Two closely *P. aeruginosa* subclades; and *H. influenzae* and *S. pneumoniae*. The OTUs are annotated with the most common keywords obtained from the metadata of their global distributions. We supply all further outlier pairs in Supplementary Tables 4&5. **c.** Violin plots depicting the community similarities of OTU-pairs which are predicted to interact based on FlashWeave (n=100, blue violin plot), the same number of randomly selected pairs (green), and random pairs corrected for phylogenetic relatedness bias (orange). Vertical line denotes the mean in each violin plot. ***=p_Mann-Whitney U_<0.0001.

Applying this observation to individual phyla (refer to Extended Data Fig. 9 and Fig. 4c), it is tempting to speculate that phyla with shallow increases in community conservatism, such as Proteobacteria and Tenericutes, could be more generalist in nature. On the other hand, phyla with specialist-like trendlines, such as Fusobacteria and several Archaeal phyla, might indeed have specialist lifestyles. In support of this, curve steepness is significantly correlated to per-phylum aggregated generalism scores (ρ= 0.46, p _spearman_ =0.048, Supplementary Table 3). In the future, these trends could inform a more fine-grained generalism score than we utilized here, able to also take smaller niches into account.

### Outliers can be ecologically informative

Most of the sufficiently sampled OTUs conform to the trends above - but it may also be interesting to look at outliers: species pairs which are closely related but dissimilar in their communities are hinting at relatively recent evolutionary pressures to change niches. Similarly, distantly related species which are similar in their communities might depend on each other or have a shared niche requirement independent of phylogeny. We provide a list of both types of outliers (Supplementary Tables 4&5) and highlight two examples in detail below (Fig. 5b).

On the bottom-left corner of the overall distribution plot are two *Pseudomonas aeruginosa* subclades, which are closely related. Yet, against global conservatism trends, they occupy different communities, hence hinting at a strong ecotype difference between both strains. To better understand their respective niches, we investigated all samples in which the OTUs are detected through metadata keyword summaries. This analysis indicates that P*. aeruginosa* clade 1 is adapted to the human host and enriched in samples of patients with cystic fibrosis, while P*. aeruginosa* clade 2 is a generalist found in many non-human environments. This overlaps with existing research showing that *P. aeruginosa* can be found in both niches ^59–61^. On the other hand, the OTU-pair of *Haemophilus influenzae* and *Streptococcus pneumoniae* are unrelated, belonging to distant phyla. Nevertheless, they share many community members and are both abundant in the human oral cavity and lungs (Fig. 5b), where they occasionally even form biofilms together ^62^.

This and other examples led us to the hypothesis that community similarity is informative when identifying ecologically interacting OTU-pairs: OTUs with more similar background communities should be more likely to interact. To investigate this, we analyzed all investigated OTU-pairs with FlashWeave, a software package that statistically predicts ecological interactions between OTUs ^20^. And indeed, OTU-pairs predicted to interact this way show much higher community similarity (p_Mann-Whitney U_<0.0001, Cohen’s d: 1.37), also when correcting for phylogenetic relatedness (p_Mann-Whitney U_<0.0001, Cohen’s d: 0.56, Fig. 5c). Together, these observations and the underlying data could prove useful to improve the inference of interacting or niche-defining OTU pairs.

## Outlook and conclusion

### Can we reintegrate ecological information into species delimitation?

The species concept in bacteria and archaea is still under debate. Some argue for a strict operational approach utilizing phylogenetic marker genes, usually by implementing a chosen species-level threshold (e.g.,97% for 16S rRNA, 96.5% for average nucleotide identity (ANI) of the whole genome)^63,64^. Others argue that this procedure is too simplistic, and that phenotypic and ecological information should be considered as well ^65^. In any case, most agree that delimitating species using specific thresholds (e.g. within the Genome Taxonomy Database) is pragmatic and operational, but not always ideal ^66^. While in some cases bacterial strains that belong to the same assigned species unit may differ strongly in their environmental role, others might be traditionally assigned to two different species, while performing the same principal role in the ecosystem.

Previous research has argued that a distribution-based approach could be used to improve OTU-delimitation ^67,68^. Here we propose to build upon these ideas, and instead of solely relying on marker similarity (Fig. 6a), to reintegrate ecological information into species delimitation in the future. We suggest to achieve this specifically using community conservatism (Fig. 6b), resulting in a combined clustering strategy (Fig. 6c). Operationally, we envision a two-step OTU clustering approach: First, a purely sequence based OTU-clustering would serve as an initial cursory analysis point, used solely to obtain pairwise community similarity values. These ecological similarities could then be integrated with pairwise genomic similarities into a new weighted metric that guides delineation of ecologically informed OTUs (eOTUs). This combined approach should yield a more natural OTU-clustering, ideally alleviating some of the disagreements between “classical”, phenotypically and ecologically informed taxonomy versus sequence similarity-based OTU clustering. The choice of how highly to weigh sequence similarity versus community similarity will be subject to empirical and theoretical considerations. Similarly, while we here used pragmatic, data-driven measures for sequence identity and community similarity, the choice of metrics is flexible. A hypothetical example with 4 OTUs, using the example metric “0.5 * 16S rRNA sequence identity + 0.5 * average BCS” is provided in Extended Data Fig. 10 and highlights how niche preferences could result in an alternative clustering outcome. We acknowledge that full implementation will require fine-tuning and benchmarking work that is outside of the scope of this article. For now, we wish to highlight that community preferences and their conservation trends are easily assessed from cross-sectional data (in contrast to other relevant phenotypes) and show promise for more ecologically meaningful species delimitation.

**Figure 6:**
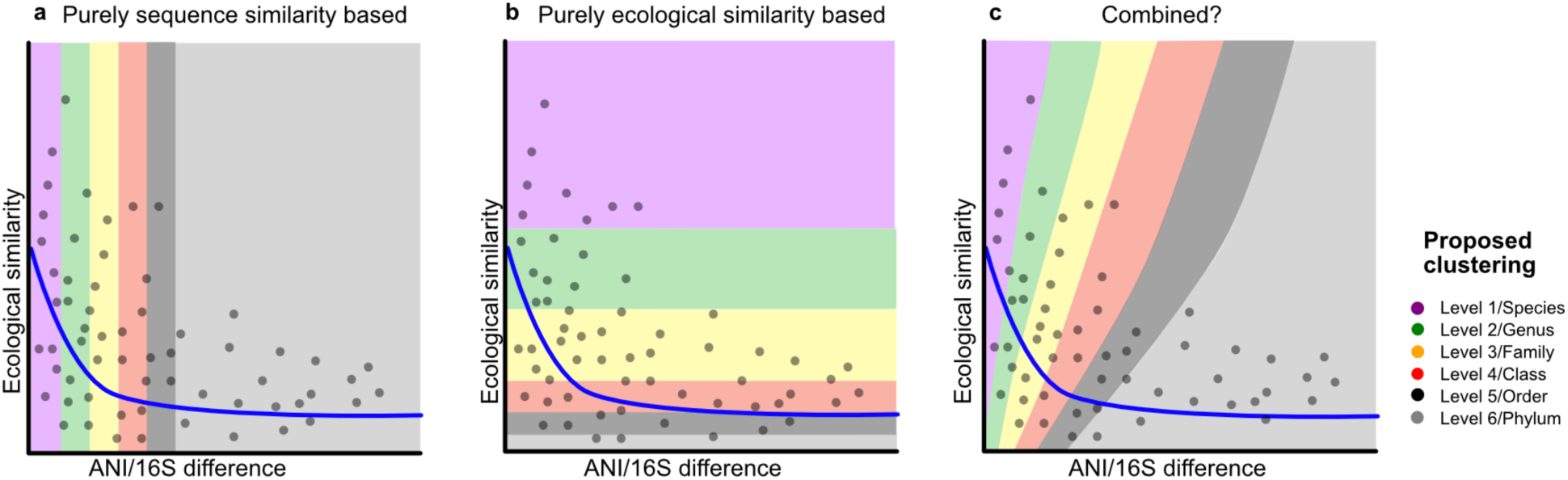
Reintegrating ecological information into species delimitation. **a.** Each grey dot denotes a hypothetical pair of Operational Taxonomic Units (OTUs). The classical assignment of species into OTUs only takes the sequence similarity into account (usually: 16S rRNA or average nucleotide identity (ANI)). **b**. A hypothetical classification of OTUs based only on ecological niche similarity. **c**. We propose a combination of both as a more realistic system: Utilizing ecological information (such as community conservatism) reintegrated into OTU-clustering, taking both sequence similarity as well as ecological information into account when forming ecologically informed OTUs: eOTUs.

## Conclusion

We found that community conservatism is universally present in all investigated phyla and environments, on a global scale. We postulate that this community conservatism signal could be useful to infer how quickly a given microbial lineage usually adapts to new environmental conditions (or communities). Potential applications include microbiome engineering, where such inferences could improve predictions of species addition or removal effects in a given community, based on their niches ^69^.

Of note, there are sampling biases associated with publicly available compositional data. For example, imbalance of sampled environments and differences in sequencing depths or protocols may influence community similarity values. However, such biases are independent of taxonomic relatedness, and thus should not generate the trends presented here.

Niche conservatism and phylogenetic signal are well-established concepts in the study of animals and plants, but their assessment in microbes has been limited by the challenges in ascertaining phenotypes and niches in free-living organisms. The concept of community conservatism offers an alternative approach to investigating these patterns in microbial communities, as well as the prospect of reintegrating ecological information into species delimitation.

## Methods

### MicrobeAtlas data retrieval

We searched the NCBI Sequence Read Archive^70^ or samples and studies containing any of the keywords ‘metagenomic’, ‘microb*’, ‘bacteria’ or ‘archaea’ in their metadata and downloaded the corresponding raw sequence data. To assign OTU labels, quality filtered data were mapped to MAPref v.2.2.1 using MAPseq v.1.0 at a ≥0.5 confidence level^35^. We then filtered out samples containing less than 1,000 reads and/or less than 20 OTUs defined at 97% 16S rRNA gene identity, and further retained only samples with at least 90% estimated community coverage ^71^.

NCBI Sequence Read Archive sample metadata were parsed to classify every sample into five general environments: animal, marine, freshwater, plant and soil. Each OTU was then also assigned a main environment, based on sample prevalence. These environmental assignments and keywords can be found in the file “samples.env.info” obtained from https://microbeatlas.org/index.html?action=download, on March 10th, 2023. The identifiers of all OTUs used in Figures (Fig. 2 and Fig. 5b) are provided in the Supplementary Information.

### Selection of OTU pairs, exclusion criteria

We used stringent criteria in selecting the OTUs (on a 99% similarity threshold) and samples which we analyzed. We only compare samples which do not belong to the same project ID. Additionally, we only consider samples which include at least 20 different OTUs and over 1,000 reads. Each OTU is only allowed in a maximum of 10 comparisons (increased to 30 in phyla <3,000 OTUs) to avoid overrepresentation of certain taxonomic groups, and we remove all eukaryotic reads to solely focus on the prokaryotic diversity.

For the general trend we compared 25,000 pairs, whereas for the phyla-specific and environment-specific points we used 10,000 pairs each. To obtain a uniform distribution of distances, we created 50 bins of phylogenetic distances and filled each bin with randomly drawn pairs within that reach. We removed the lowest 3% of distances, since they may contain some misclassified OTUs, or cases where the bin would otherwise be impossible to fill. Each OTU-pair was only selected once. The taxonomy of the OTUs was assigned according to the NCBI assignments of the representative 16S rRNA sequences.

### Phylogenetic tree generation

Phylogenetic Trees were generated from the alignments of reference sequences in MAPref v.2.2.1 using fastTree 2.1.10 with the “-nt -gtr -gamma” parameters ^72^, and multifurcations were removed subsequently. Phylum-specific trees for each phylum with >500 OTUs were generated with the same evolutionary model to ensure comparability. Tree distances were extracted using the distance function of ete3 version 3.1.2 ^73^.

### Fraction of shared samples and sequence similarity

For all OTU-pairs which were compared in their tree distances and community similarities, we additionally calculated the sequence similarities of the full-length representative 16S rRNA sequences with a custom script. We furthermore calculated the fraction of shared samples based on the overlap in prevalence within MicrobeAtlas.

### Calculation of community conservatism

The Bray Curtis similarity (also called Quantitative Sørensen–Dice index) was calculated with the formula (1 - Bray Curtis dissimilarity). HPC-CLUST v1.1.0 ^38^ was used for the calculations with the following parameters: -“t samples -nthreads 30 -dfunc braycurtis_skipproj -makecluststats -projf”. For each OTU-pair, we compared all samples which do not belong to the same research project (i.e., do not share the same “project-ID” at the Sequence Read Archive) in which they are detected in a pairwise manner (e.g. if OTU 99_1 if found in samples A and B, and OTU 99_2 is detected in samples A, C and D we would compare the community similarities of A-C, A-D, B-C and B-D. A-A would not be compared). We record multiple quantiles but use the mean in all plots unless specified otherwise. The output was further processed with pandas v1.0.3 ^74^ and plotted with bokeh version 2.2.3. We used a locally weighted scatterplot smoothing (lowess, statsmodel.api.nonparametric.lowess, frac=1/5) as well as an exponential decay function (scipy.optimize.curve.fit) to fit the data. Final plots were adjusted in affinity designer. All custom code is available on github under https://github.com/lukasmalfi/community_conservatism.

### Null model generation

To create a general null model, we randomly compared the communities of 50,000 randomly chosen samples with the same parameters as described above. We furthermore created individual baselines for all environmental combinations (so, only comparing soil-soil samples, animal-animal, animal-soil etc.). We then used these values (Supplementary Table 1) to generate phylum-specific baselines. To achieve this, we used the assigned main environment of each OTU and computed the ratios of their environment combinations (e.g.: animal-animal: 0.1, animal-soil: 0.05, animal-aquatic:0.03 etc.) and then calculated the respective null model. To plot the phylum-specific trendlines, we normalized those and divided the mean community similarity models by the calculated null-model.

### Phylum specific ratios

Since many of the OTUs are not taxonomically annotated to species or genus level, we estimated the approximate range of species and genus-OTU pairs from the general trend. We used the middle 60% of rank-specific distributions (i.e. excluding the top and bottom 20%, respectively) to obtain “species-level”, “genus-level” and “phylum-level” bins based on the phylogenetic distance. We then calculated the average community similarities of those 3 bins for each phylum. As a next step, we divided each species-level bin by the other two to create the ratios used to estimate the increase in community similarity from the genus level to species level, and from the phylum baseline to the species level.

### Outlier OTU-pairs

We classified OTU-pairs as outliers on both extrema: i) pairs which are very closely related (tree branch length < 0.2), yet very different in their communities (mean Bray Curtis Similarity < 0.04), and ii) pairs which are not related (tree branch length > 0.8), yet their communities are similar (mean Bray Curtis Similarity > 0.08). Additionally, we only considered outliers for which at least 10,000 sample comparisons had been calculated. We provide a list of all outliers that fall into these bounds in Supplementary Tables 4 & 5.

### Generalist and specialist analysis

We calculated an ‘environmental flexibility’ index for each OTU based on its abundance distribution across animal, aquatic, soil, and plant environments. OTUs with more uniform abundances across these environments (measured as Shannon entropy) scored higher on the index, indicating greater generalism. Conversely, OTUs with uneven abundances, showing preference for specific environments, scored lower - suggesting more specialized adaptations. In Fig. 4a we plotted 10,000 “Specialist” OTU-pairs (lowest environmental flexibility score) and 10,000 “Generalist” OTU-pairs (highest environmental flexibility score). The individual generalism scores of all 99% OTUs with taxonomic annotations were aggregated to obtain phylum-level generalism scores. These were correlated to the increase of community conservatism from species to genus level (ratios) with a Spearman correlation using the stats.spearmanr function of the scipy package v1.4.1.

### Word Clouds

For each OTU, keywords of all samples in which they were found were added to a list using custom code in Python 3.7.6. The list of obtained keywords was used to create a word cloud with WordCloud v1.5.0^75^ with a custom color map and the following parameters: stopwords = stopwords, prefer_horizontal = 1, min_font_size = 10, max_font_size = 150, relative_scaling = 0.4, width = 1000, collocations = False, height = 400, max_words = 15, random_state = 1, background_color = “white”.

All custom code, as well as the used stopwords are on github.

### Interaction network analysis

We analyzed the OTU pairs plotted in Fig. 3 by constructing a global network of predicted interactions. We employed the local-to-global learning approach, as described by ref^76^, using FlashWeave v.0.19.0 ^20^. This method generates a Bayesian network skeleton, representing potential ecological relationships between species while accounting for ecological or technical confounding factors.

FlashWeave’s algorithm operates in two main steps: First, it heuristically identifies likely confounding variables for each species pair based on univariate associations and previous algorithm iterations. Second, it tests whether the focal association persists when conditioned on these candidate confounders.

We configured FlashWeave with the following parameters: sensitive = false, heterogeneous = true, and max_k = 3. With these settings, the software converts non-zero read counts to centred log-ratio-transformed values, addressing compositionality issues, and then discretizes these values. Conditional mutual information tests are subsequently performed on the discretized data.

We chose the 100 OTU-pairs with the highest predicted interaction score to compare them against a random selection of 100 random OTU-pairs from the same dataset. Additionally, a second control group was chosen with a phylogenetic distribution matching the high-interaction pairs, to correct for phylogenetic relatedness. To this end, for each OTU-pair selected, a random control within +-0.025 tree branch length was drawn.

### Statistics

The comparisons of the community similarity values of different taxonomic groups were performed using a two-sided Mann-Whitney U test in the scipy package v1.4.1 (“stats.mannwhitneyu”)^77^. We calculated the differences between the interacting pairs and the control groups using a two-sided Mann-Whitney U test. Resulting P values were corrected for multiple testing using the Benjamini–Hochberg method. Effect size was calculated using Cohen’s d.

## Data Availability

All data will be made available in Zenodo. For this study, we used an older version of MicrobeAtlas which can be downloaded in Zenodo.

## Code Availability

All custom code used in the analysis can be obtained from github under https://github.com/lukasmalfi/community_conservatism.

## Contributions

L.M. and C.v.M. conceived and designed the study. L.M., J.T. and J.F.M.R generated the data. L.M. performed the statistical analyses and generated the visualizations. C.v.M. supervised the study. L.M. wrote the first draft of the article, with input from J.T. and C.v.M. All authors contributed to revising and editing the final article.

## Acknowledgements

We thank members of the von Mering lab, as well as Maarten Langen, Julien Massoni and Chiara Rickenbach for their input and helpful discussions. The work was funded by the Swiss National Science Foundation (project grant 310030_192569, as well as through one of their National Centers of Competence in Research, “Microbiomes”, 310030_192567).

## Extended Data

**Extended Data Fig. 1:**
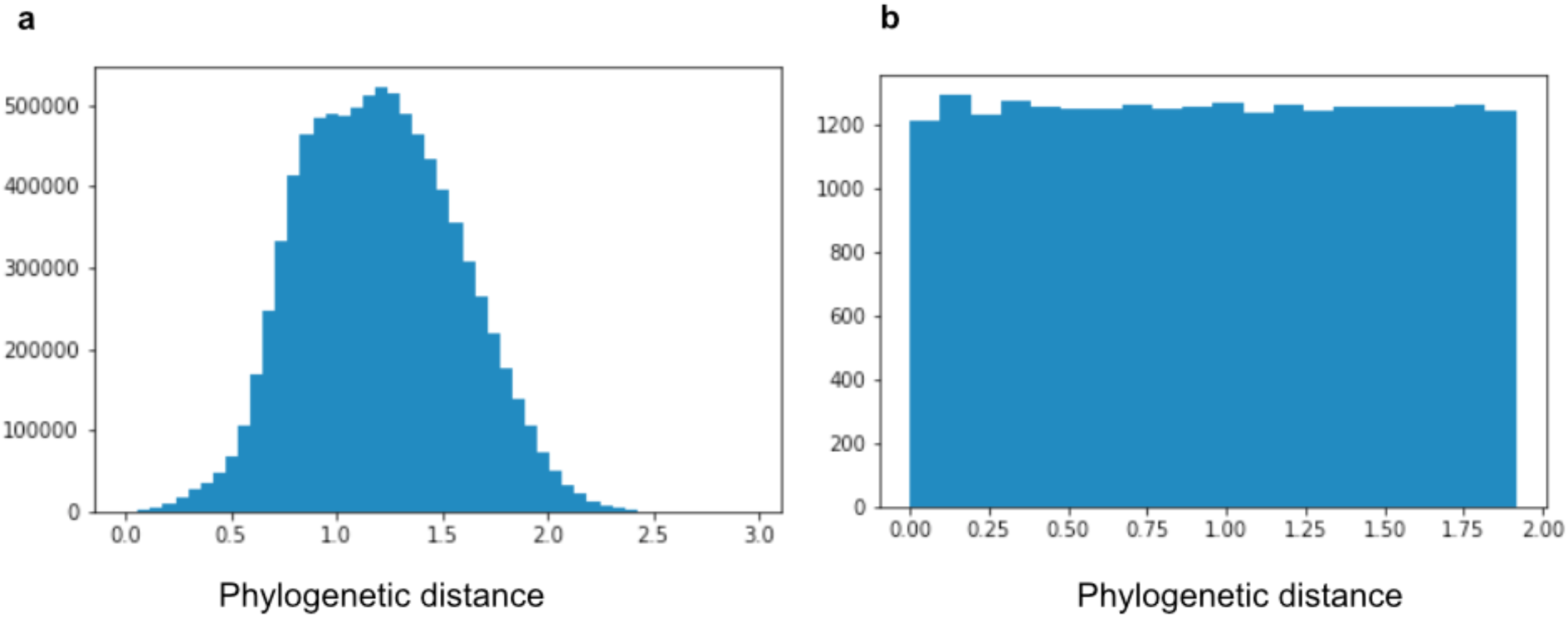
Obtaining a uniform distribution of phylogenetic distances. **a.** Downsampling the phylogenetic distances of OTU-pairs from the estimated original distribution to **b**. achieve a uniform distribution of phylogenetic distances over the whole range of relatedness.

**Extended Data Fig. 2:**
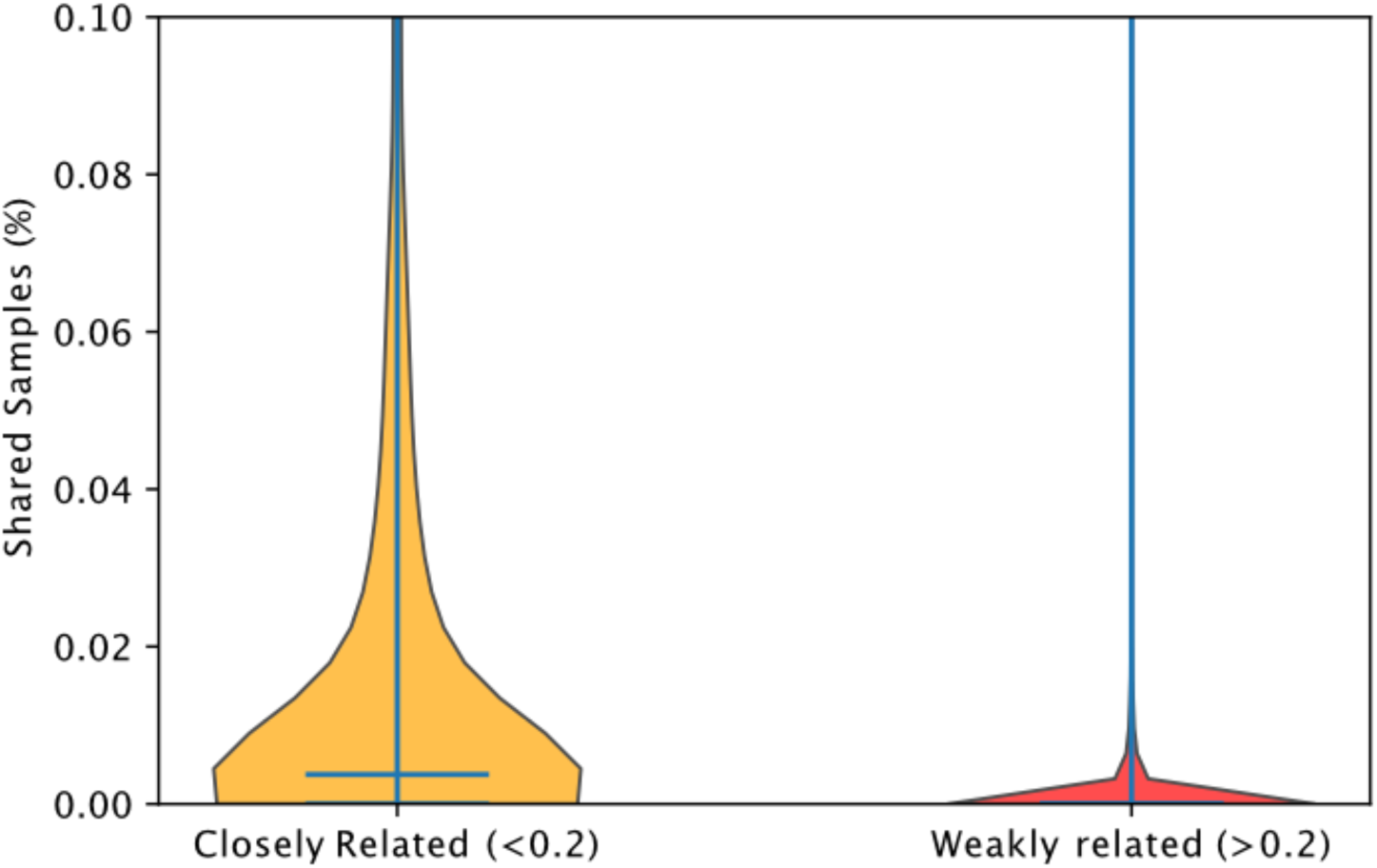
Closely related taxa co-occur in more samples. The percentage of shared samples in closely related OTUs (phylogenetic tree branch length <= 0.2) is shown on the left-hand side in yellow. Weakly related OTU-pairs (phylogenetic tree branch length > 0.2) are on the right side in red. The horizontal line denotes the mean percentage of shared samples.

**Extended Data Fig. 3:**
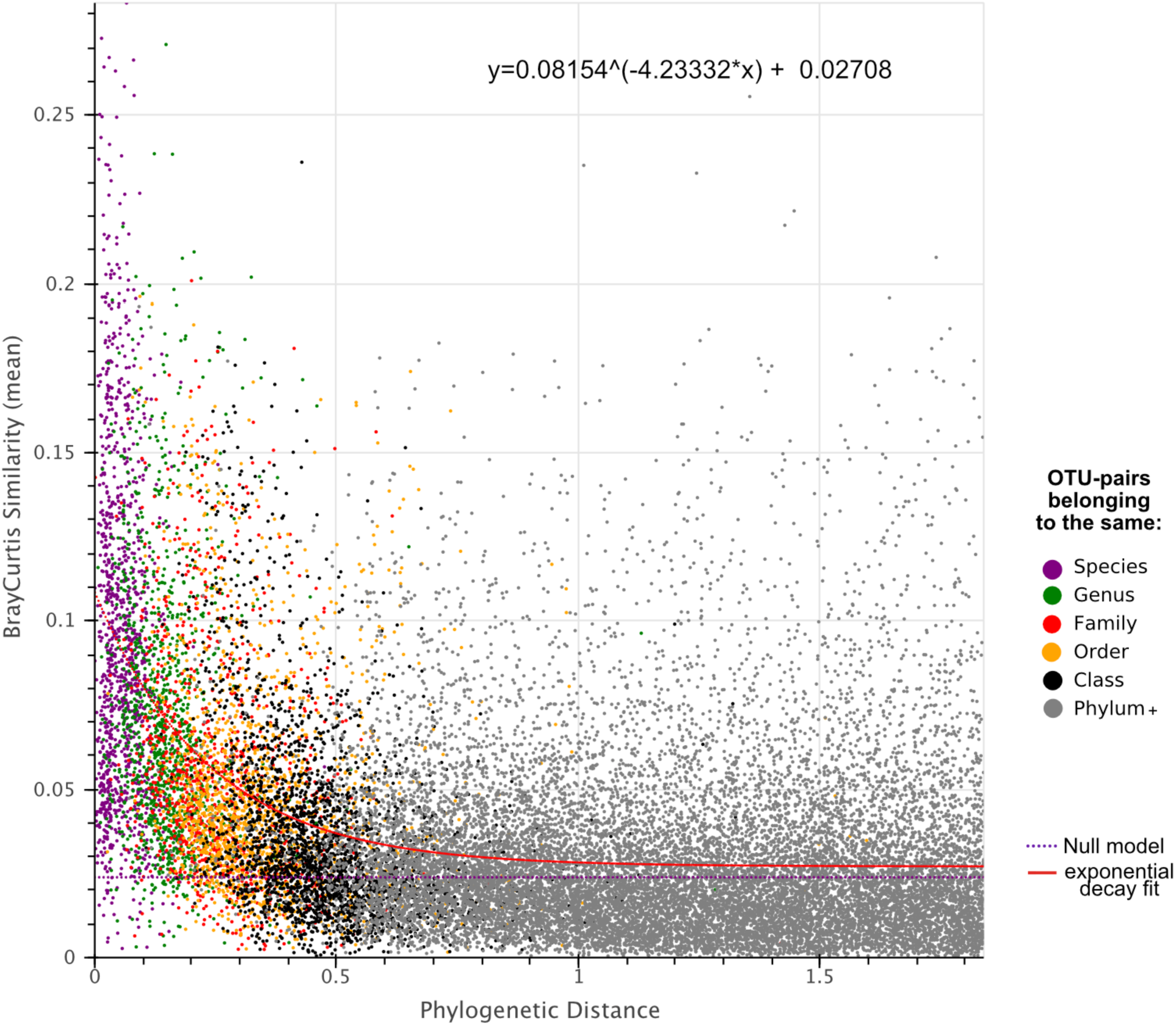
Exponential decay fit and function. Community similarity falls as phylogenetic distance increases, visualized here through 25,000 OTU pairs with available taxonomic annotation to species level: exponential fit (and formula) and random expectation are shown as blue and red lines, respectively. Each dot corresponds to one OTU-pair colored according to their most specific shared taxonomic rank, with their relatedness shown on the x-axis and the average similarity of their communities (Bray-Curtis similarity) on the y-axis.

**Extended Data Fig. 4:**
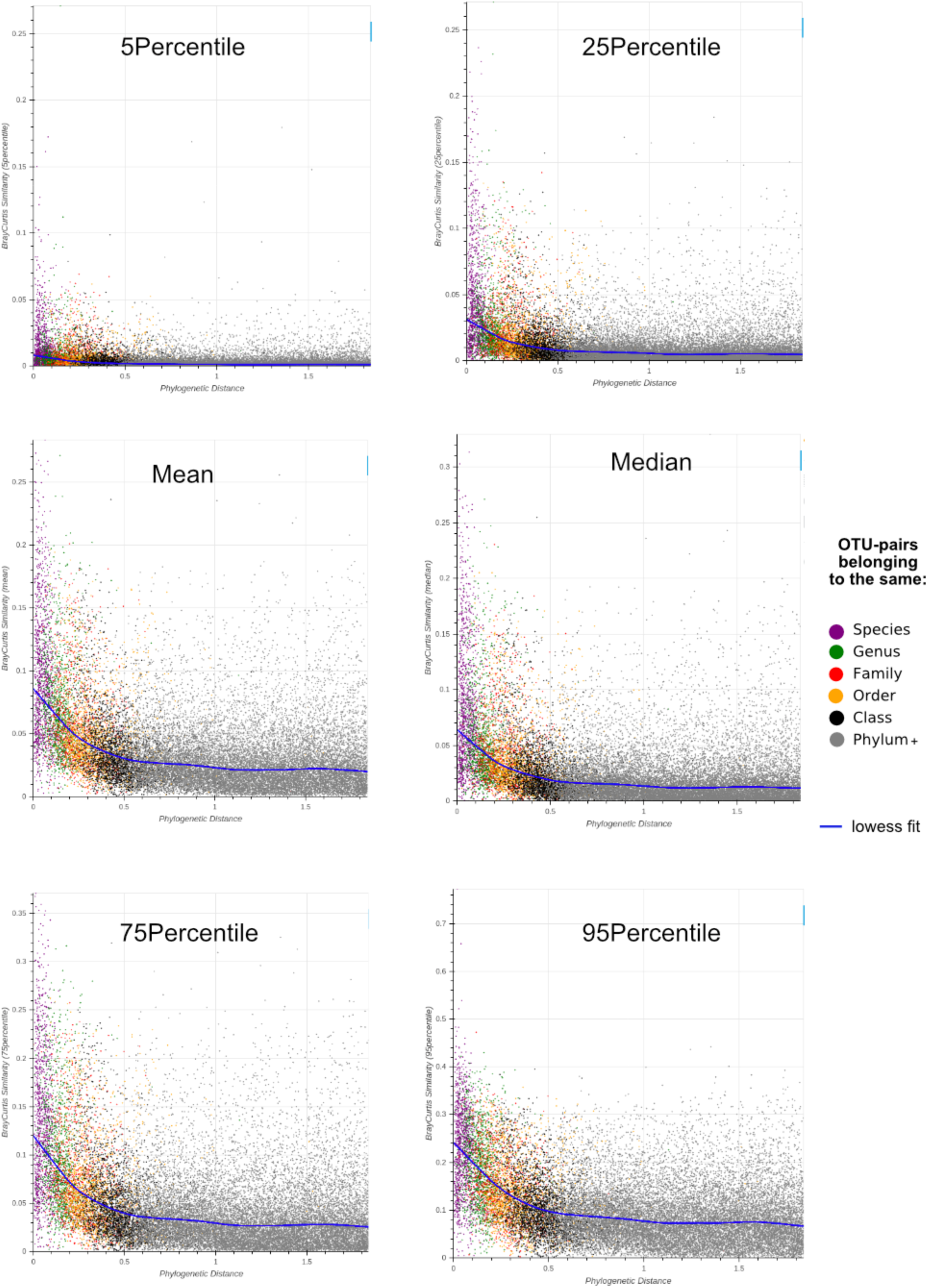
Different percentiles in community conservatism calculation. Different percentiles of community similarities are shown here, in addition to the respective lowess fit as blue lines. Each dot corresponds to one OTU-pair colored according to their most specific shared taxonomic rank, with their relatedness shown on the x-axis and the average similarity of their communities (Bray-Curtis similarity) on the y-axis.

**Extended Data Fig. 5:**
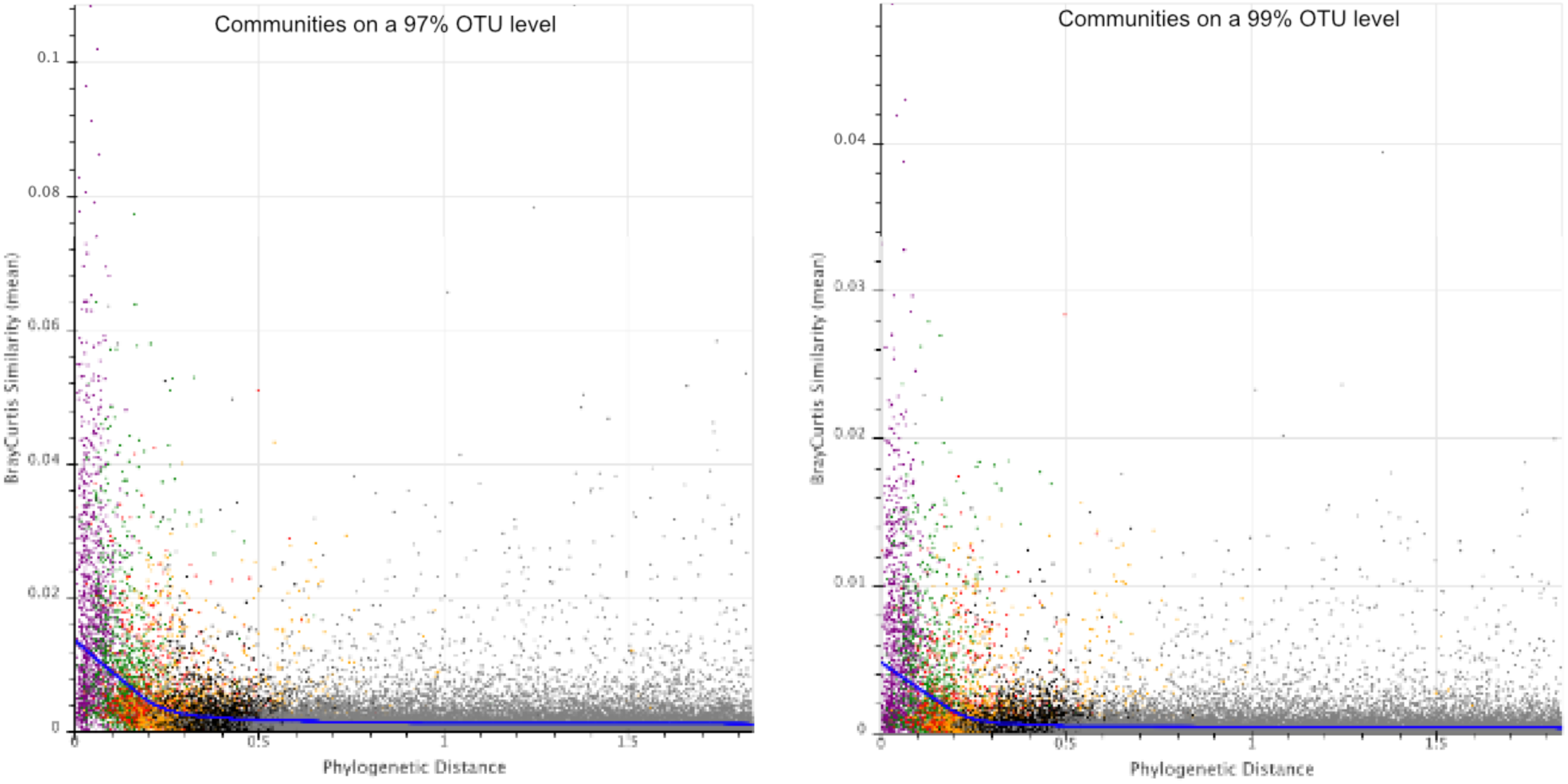
Different OTU-thresholds in community similarities. Different granularity OTUs (97% and 99% instead of 90% as in Figure 3) used to compute the ß-diversity in the microbial communities (y-axis). Each dot corresponds to one OTU-pair colored according to their most specific shared taxonomic rank as described previously.

**Extended Data Fig. 6:**
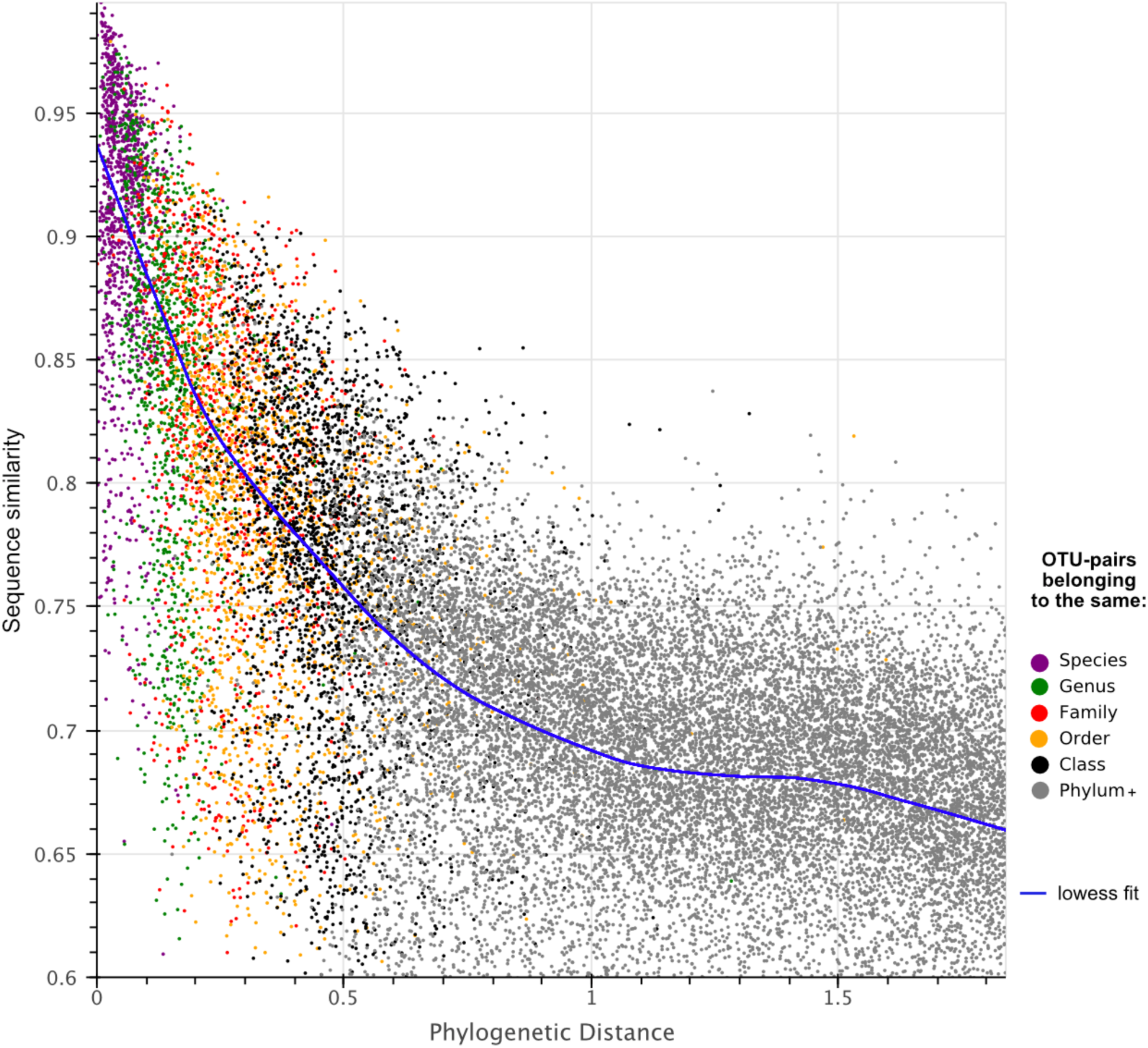
Sequence similarity correlates strongly with phylogenetic distance. The sequence similarity of the full-length 16s rRNA representative sequence of the OTU-pairs (y-axis) and the distance estimated by the tree branch length from the phylogenetic trees (x-axis) are plotted against one another. A lowess fit is applied to the data and shown with a blue line. Each dot corresponds to one OTU-pair colored according to their most specific shared taxonomic rank, with their relatedness shown on the x-axis and the average similarity of their communities (Bray-Curtis similarity) on the y-axis.

**Extended Data Fig. 7:**
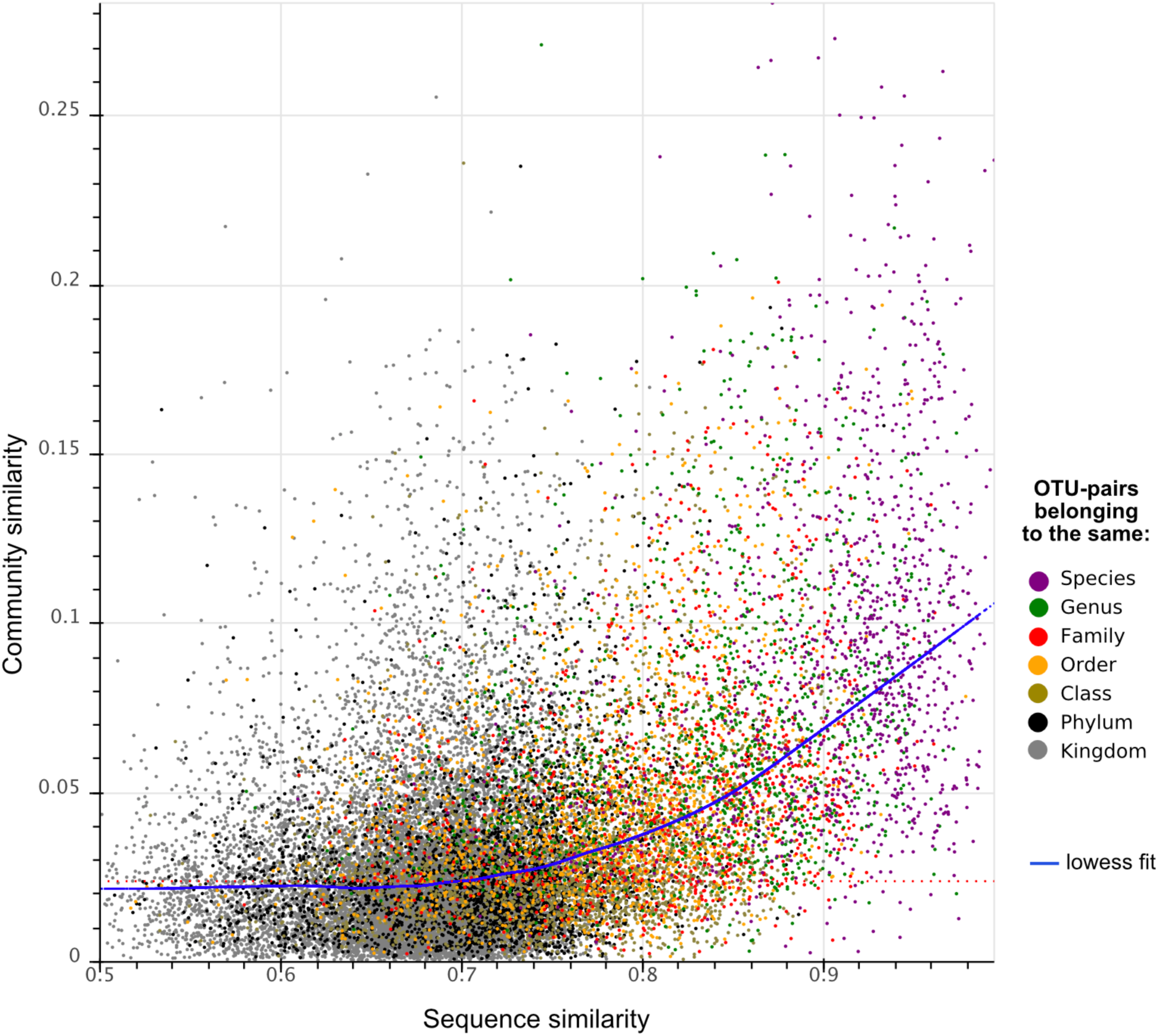
Sequence similarity offers less resolution than tree branch length. The same data as in Fig. 3, just using sequence similarity instead of phylogenetic distance as a measure of relatedness. The disagreements between taxonomic assignments and sequence similarity become apparent here. They highlight the differences between taxonomic classifications often established decades ago with the help of morphological and metabolic criteria, versus contemporary OTU clustering methods solely relying on sequence similarity. ^78^

**Extended Data Fig. 8:**
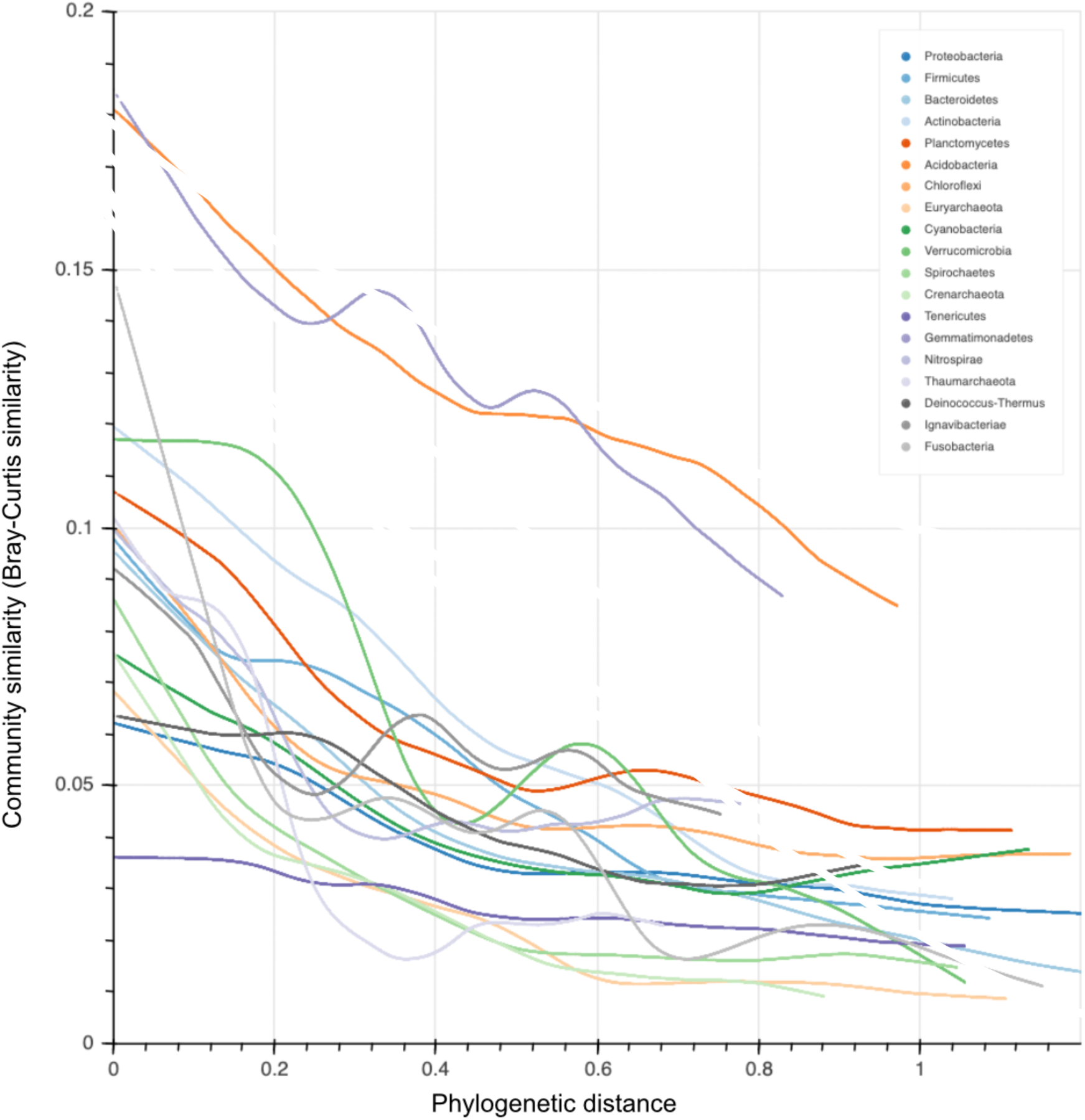
Unnormalized phyla trendlines. Lowess fits of all phyla with >500 OTUs are shown here. Each lowess fit stems from 7,500 OTU-pairs, whereas phyla with a low number of OTUs (<3,000) show an increased level of noise.

**Extended Data Fig. 9:**
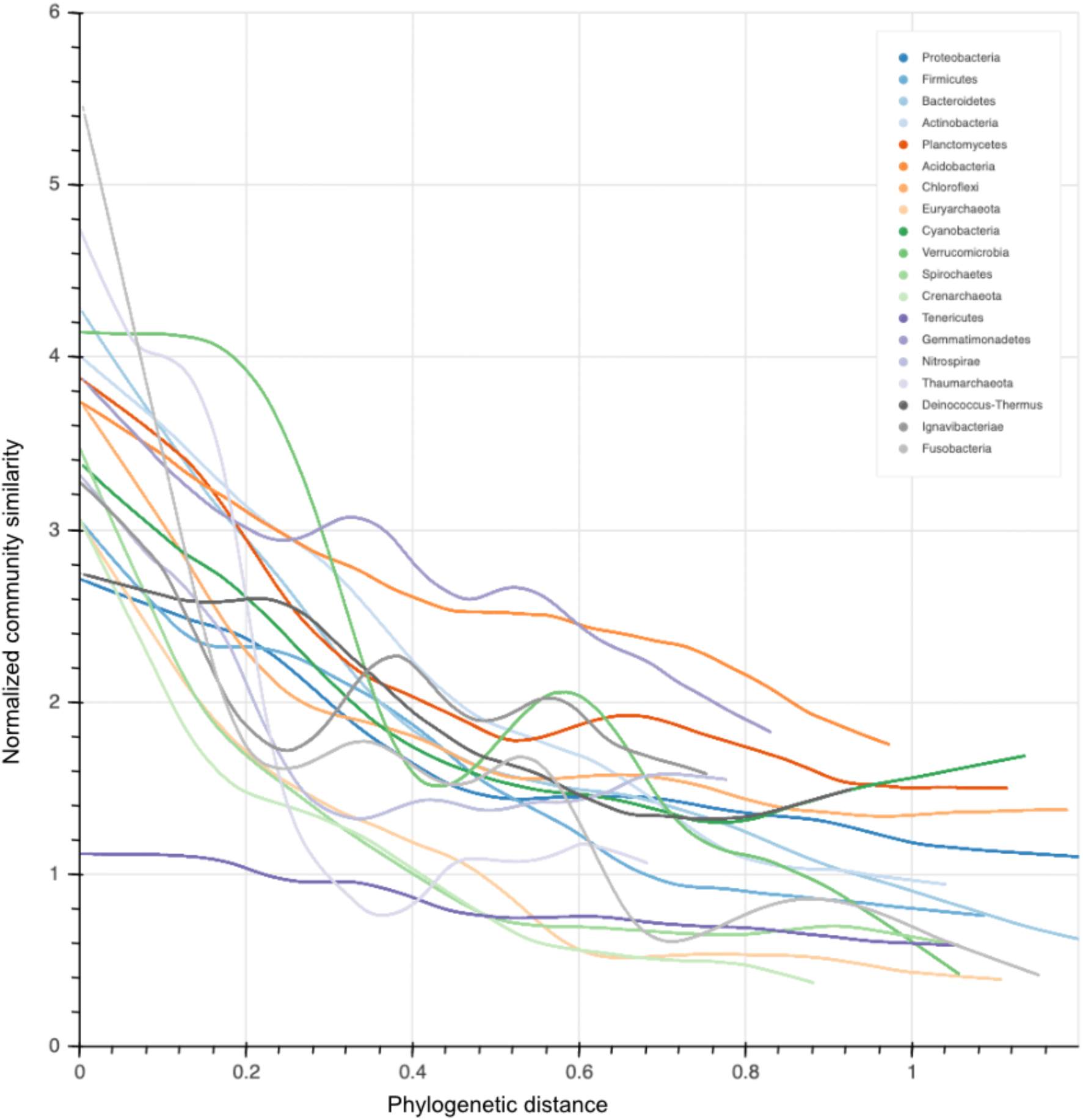
Normalized phyla trendlines. Lowess trendlines of all phyla with >500 OTUs are shown here. Each lowess fit is calculated from 7,500 OTU-pairs each, whereas phyla with a low number of OTUs (<3,000) show an increased level of noise. Each phylum is separately normalized according to Supplementary Table 1 (See Methods).

**Extended Data Fig. 10:**
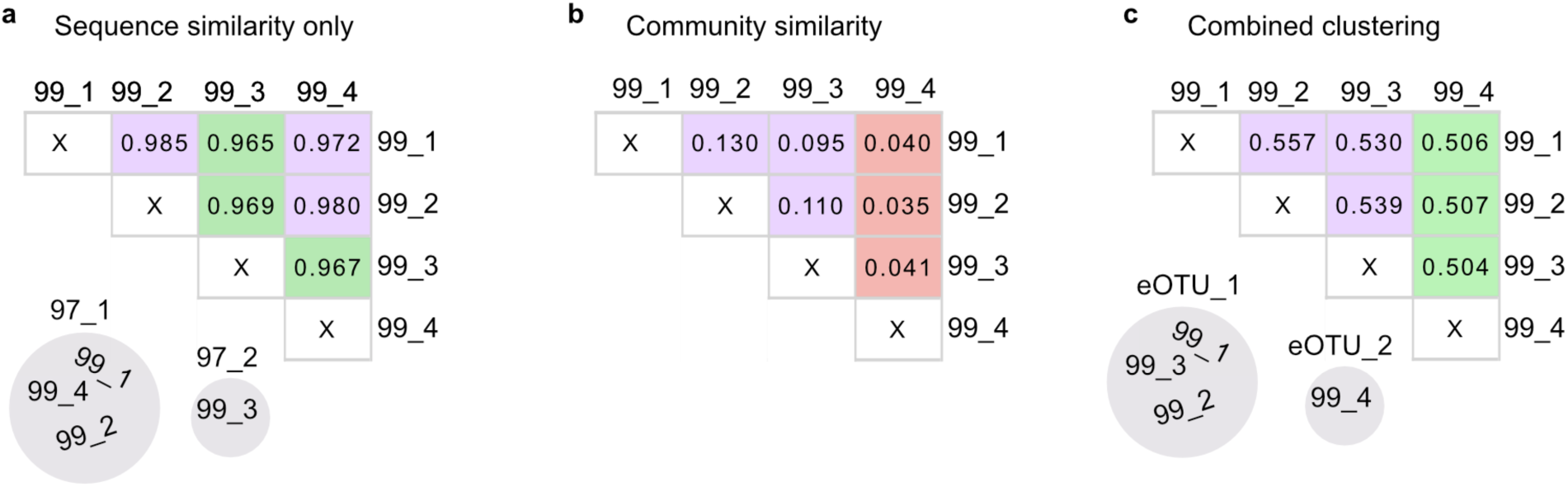
Hypothetical eOTU-clustering with 4 examples. In this example, four 99% OTUs are closely related. **a.** Pair-wise sequence similarity values are shown in the table. When using a 97% clustering threshold in OTUs, 99_1, 99_2 and 99_4 would cluster together into one 97% (purple, species-level) OTU; and 99_3 would form a separate OTU (green). **b.** Pair-wise Bray-Curtis similarity (BCS) values are shown in the table. When investigating ecological information, it becomes apparent that 99_1, 99_2 and 99_3 are very similar in their niches (purple), whereas 99_4 appears to occupy a different niche (red). **c.** We propose to join both metrics (using the formula (0.5* 16S rRNA sequence similarity) +0.5*(BCS) for illustration) to inform the definition of ecological OTUs: eOTUS. In this hypothetical example, a value of 0.525 would delimit the four 99% OTUs into two ecologically consistent eOTUs. More specifically, considering the environmental information would result in an alternative clustering that groups the environmentally similar OTUs 99_1, 99_2 and 99_3 into one eOTU (purple).

